# Measure Catabolism: Real-time shifts in microbial metabolism through online pressure measurements

**DOI:** 10.64898/2026.06.08.729509

**Authors:** Martin Malthe Borch, Ugochi Jennifer Nwaokorie, Philip J. Gorter de Vries, Kaspar Valgepea, Alex Toftgaard Nielsen

## Abstract

Microbial activity is often inferred from cell density measurements; however, biomass formation is merely an indirect, cumulative result of metabolism, known as anabolism. Microbial activity is more accurately indicated by energy conservation or catabolism. This is especially true under low or no-growth conditions, where anabolism remains constant, and shifts in catabolic fluxes go unnoticed with biomass measurements alone. In anaerobic and gas-based metabolic processes, net gas exchange is linked to energy conservation, and catabolism can then be quantified through headspace measurements.

We introduce a sealed-vial, non-invasive workflow that uses high-resolution headspace pressure measurements to estimate gas exchange rates and catabolic reactions, enabling real-time visualisation of metabolic shifts throughout an entire batch cultivation cycle. The method was applied to carbon monoxide (CO) fermentations of three *Clostridium autoethanogenum* strains (JA1-1, LAbrini, and LAbrini_mut) cultivated in serum bottles. Two of them were indistinguishable by OD-derived μ_max_. Pressure-derived gas uptake rates resolved multiple exponential phases of gas consumption and identified specific shifts in the metabolism, consistent with transitions from mixotrophic to autotrophic growth. Small but significant differences in terminal headspace pressure were detected, providing an experimentally accessible end-state parameter for phenotypic characterisation that would be obscured by routine intrusive headspace sampling. Finally, pressure-derived catabolic rates further enabled estimates of relative product formation during the main autotrophic phase.

The strains were successfully characterised and distinguished by identifying several exponential phases of gas consumption and their rates, as well as differences in the final absolute pressure threshold. This provided phenotypic characterisation and insights not obtainable from OD measurements alone. The work establishes a practical framework for catabolism-resolved microbial characterisation in sealed batch vials through high-resolution online pressure (gas exchange). The assumption that pressure measurements correlate with CO_2_ and catabolic rates is sensitive to solubility/buffering and temperature/vapour effects, but these limitations are addressable through controls and complementary analytics.

## 1 Introduction

Microbial activity is historically measured through the growth of biomass and an increase in turbidity (Pasteur 1857), but biomass formation is only one part of the microbial activity and the indirect, cumulative result of metabolism, referred to as anabolism. Total microbial activity is more accurately indicated by energy conservation or catabolism. This is especially true under low or no-growth conditions, where anabolism remains constant, and shifts in catabolic fluxes go unnoticed with biomass measurements alone. In industrial anaerobic biotechnological processes, the products are mainly formed during catabolic reactions. Examples of common, widespread anaerobic biotechnological processes include bioethanol and biogas production. In anaerobic metabolic processes, the catabolic energy conservation through redox reactions often involves either the production or consumption of gases, while anabolism contributes less to the overall gas exchanges (Table 1). Pressure is a physical observable parameter, just like OD, and it enables quantification of catabolic rates from headspace pressure measurements in closed anaerobic cultivations. Pressure sensing thus enables measurements of the catabolic reactions and can reveal metabolic shifts, also when biomass formation cannot be measured or does not change.

**Table 1.**
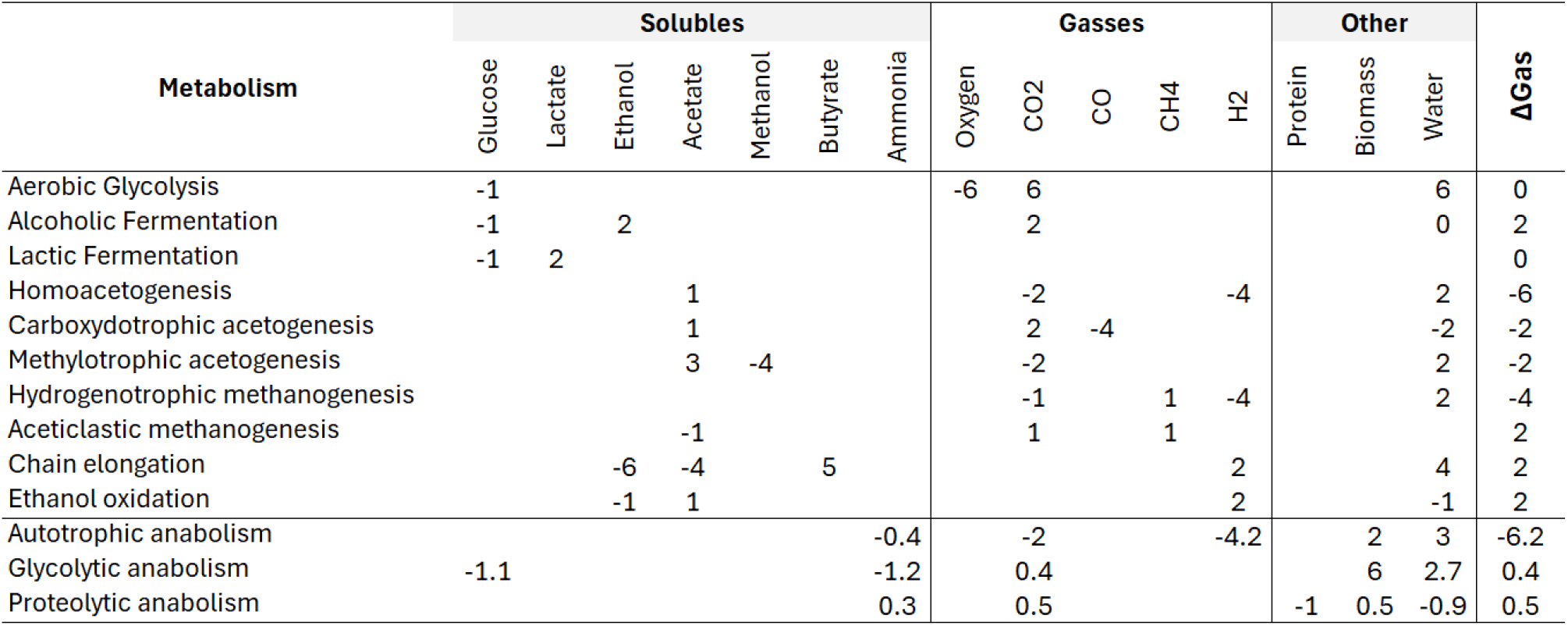
catabolic reaction stoichiometry. Calculated as outlined in (Shuler and Kargi 2002) chapter 7, c-mol biomass approximation of: CH_2_O0_.5_N_0.2_. Δgas is the sum of all stoichiometric coefficients of gaseous substrates and products. In the table, substrates are negative values and products positive. Elemental composition of protein is the average elemental composition of amino acids, ignoring non CHON elements.

In gas-fermentation, the gases such as CO_2_, H_2_, and CO are consumed, with applications in the production of sustainable chemicals and biomanufactured materials (Redl et al. 2017). In small-scale laboratory autotrophic cultures, pressure, OD, and product off-line analysis (HPLC) are routinely monitored through manual sampling. This disturbs anaerobic cultures through leaks, contamination risks, and transient temperature effects, and manual sampling makes reliable rate estimation near-infeasible (Borch, Kehr, et al. 2026). In larger bioreactors, sampling can be complemented by online gas mass spectrometry (MS), which monitors the formation of volatile and gaseous products in real time and provides insights into catabolic reactions. Online product analysis using MS, though, requires a constant gas flow to transport the products to the analyser. For pure-culture autotrophic cultivations, gas sparging is also used to supply gaseous substrates. But, sparging is less suited if the desire is to analyse anaerobic microbial community interactions. In microbial communities, reactions can occur in cascades within microbial networks. Here, a product from one organism becomes the substrate for the next reaction in another microorganism. As an example, dark fermentation of sugar substrates produces H_2_/CO_2_, which can then be converted to acetate by acetogens (Borch, Grimalt-Alemany, et al. 2026). But the interaction can occur and be observed only if the intermediate products are not washed out by sparging the reactor with N2 or another carrier gas used for an online gas analyser, both of which are common practices. Another advantage of undisturbed, high-quality online pressure measurement in closed-batch cultivations is the detection of gas thresholds. This is the point at which the energy driving (thermodynamic) reactions is balanced, and catabolic processes stop. This can be used to measure the energy generated, ATP mol/product (Laura and Jo 2023; Philips 2020).

As mentioned in anaerobic processes, product formation is linked to catabolism and gas rates. This enables estimation of the product rate by subtracting the biomass growth rate from the overall pressure change rate, as outlined below. While this is a useful screening approach, it should not be interpreted as a quantitative molar product rate without additional stoichiometric constraints and gas composition data. If the gas composition is not known, the method can still be applied if the catabolic reactions are known or controlled by the experimental design.

The gas ratio between substrates and products of the catabolic reactions varies widely (Table 1). Some examples are respiration, endogenous metabolism with production and consumption of storage compounds, and hydrogenotrophic methanogenesis (Table 1). When the cells in a pure culture or a microbial community change catabolic reactions, this can be observed independently of changes in the total biomass pool. To summarise, applying pressure measurements in closed-batch cultivations addresses the challenges of manual pressure or OD sampling and of gas sparging in bioreactors. The key points are:

- Pressure measurements do not require growth to measure microbial activity.
- Rate estimates can be obtained through simple single-batch experiments without requiring intrusive sampling, complex chemical analysis equipment, or a chemostat experiment.
- Sparging is not applied, and headspace gases are not exchanged or disturbed, allowing analysis of sequential microbial process cascades.
- In batch cultivation, microbes experience the same conditions simultaneously, leading to parallel growth. When a threshold is reached, the entire population shifts its metabolism at once. This creates a clear signal, enabling the detection of catabolic changes regardless of biomass measurements.
- The online pressure measurements enable disturbance-free cultivation, avoiding headspace disturbances and thus providing higher resolution and more accurate data.
- Furthermore, online backscatter measurements could be combined with online pressure measurements. This would enable the simultaneous quantification of catabolic and anabolic rates in a single batch experiment. This approach can be used to determine maintenance variables and track metabolic shifts throughout an entire growth cycle, as described by (Benthin et al. 1994).

In this study, we examined the relationship between headspace pressure and catabolism. Uninterrupted online pressure measurements during closed-batch cultivations were used to quantify catabolic rates and supplement existing OD analyses for phenotypic strain characterisation. Recently developed online pressure sensors (Borch, Kehr, et al. 2026) were used. We characterised three *Clostridium autoethanogenum* strains during carbon monoxide cultivation, two of which had identical growth rates, as determined by manual OD measurements. We explored the catabolism-focused characterisation approach and confirmed the broader potential of the method and the sensors. Several shifts in the catabolic rates and pressure-based parameters were identified, enabling the detection of phenotypic differences.

## 2 Materials and Methods

### 2.1 Autotrophic batch cultivation of *C*.*autoethanogenum* strains

Autotrophic batch cultivation of *C. autoethanogenum* strains was performed in chemically defined PETC-MES liquid medium (Nwaokorie et al., 2023) supplemented with 1.5 g/L of yeast extract and 0.4 g/L of cysteine-HCl·H_2_O as a reducing agent. The medium was prepared anaerobically, and the pH was adjusted to 5.7. Batch fermentations for each strain were performed in 125 mL serum bottles containing 25 mL of liquid. Six biological replicates were incubated horizontally at 37°C with orbital shaking at 120 RPM under strictly anaerobic conditions. Before inoculation, the culture bottle headspace was pressurised to 2400 mbar absolute pressure with 60% CO and 40% Ar gas mixture (AS Linde Gas). Maximal specific growth rate (µ_max_) and metabolite production data for *C. autoethanogenum* JA1-1 and LAbrini from the exponential growth phase are from (Ingelman et al. 2024), while data for LAbrini_mut strain were determined in this work using the same methodology.

### 2.2 Pressure sensor system

The pressure sensors used were the Laerke.eu pressure sensor, recently developed (Borch, Kehr, et al. 2026). The sensor measures absolute pressure in mbar from 0 to 30 bar.

### 2.3 Statistics and replication

Error bars represent variability across biological replicates and are reported as standard deviation. When pressure and OD/backscatter were measured in separate experiments, we treated cross-comparisons as qualitative.

## 3 Results

The aim was to characterise closely related *Clostridium autoethanogenum* strains and differentiate them phenotypically using pressure sensors to identify growth and production advantages resulting from strain engineering. The strains were previously characterised through OD measurements, and two of the strains could not be separated based on µ_max_ alone. Online pressure measurements of undisturbed cultures were applied for additional insights into the catabolism of the strains. The analysis revealed different gas exchange patterns and thus metabolic differences that was not observed by OD alone, enabling a phenotypic separation of the strains on several parameters. The strains were the wild-type JA1-1 (Abrini et al. 1994), LAbrini a strain derived from a laboratory evolution effort (Ingelman et al. 2024), and LAbrini_mut a mutant variant of LAbrini (created by co-workers at University of Tartu). The cultivations were performed as previously described (Ingelman et al., 2024) in 125 mL serum bottles with 25 mL medium and carbon monoxide (CO). The bottles were not sampled during the cultivation to limit disturbance of the culture and noise in the pressure data.

We first compared the raw gas pressure curves (Figure 1B). LAbrini and LAbrini_mut had very similar pressure curves, but they reached different final pressures. We then analysed the slopes of the gas curves to verify the gas transfer capability of the experimental setup. For LAbrini, the gas transfer rate gradually increased up to hour 38, after which it dropped sharply. This indicates high catabolic activity up to that time, possibly associated with the exponential growth phase. No linear gas-transfer period was observed in the slope plot, showing that the experimental setup provided sufficient gas transfer; however, after 38 hours, suboptimal metabolism is evident. A similar analysis was performed for LAbrini_mut and JA1-1, which increased gas transfer until h 40 and 90, respectively (Supplementary Figure 2). The pressure change was then plotted, normalised to the maximum pressure reached. This provides an “OD-familiar” visual representation of increasing gas consumption (Figure 1C). We identified a significant difference in total gas consumption. Log-transformation of the “gas consumption” shows the exponential gas-consumption phases as straight lines (Figure 1F). As the overall gas rate is a function of both the growth and non-growth associated maintenance, an exponential increase in pressure can be assumed as a result of catabolism coupled to an exponential phase of self-replication. For LAbrini, this is confirmed as an exponential growth phase up to hour 38 (Figure 1A), as also indicated from the gas-transfer slope scatter plot. We identify two clear exponential phases for LAbrini and three for LAbrini_mut and JA1-1 (Figure 1F), and Supplementary Figures 2, 3, and 4). All three strains exhibit a higher gas-consumption rate during the first 20 to 25 hours. We assume that these higher initial gas consumption rates are due to mixotrophic growth (the medium contains yeast extract, sodium acetate, and cysteine). For LAbrini_mut, we hypothesise that the initial two phases are both mixotrophic. During the exponential mixotrophic gas-consumption phase of LAbrini and JA1.1, they both take up gases faster than during pure-gas exponential growth. This is similar to LAbrini_mut, but here no clear shift is observed, and the rate gradually declines to the exponential gas-phase rate. Interestingly, the gas consumption rates during the late mixotrophic phases are quite similar. JA1-1 has two different gas consumption rates during the autotrophic phases at approx. 34-44h and 53-67h (r^2^ > 0.99 for both phases). Further experiments are needed to clarify the causes of these different gas-rate phases and the associated metabolic shifts observed.

**Figure 1.**
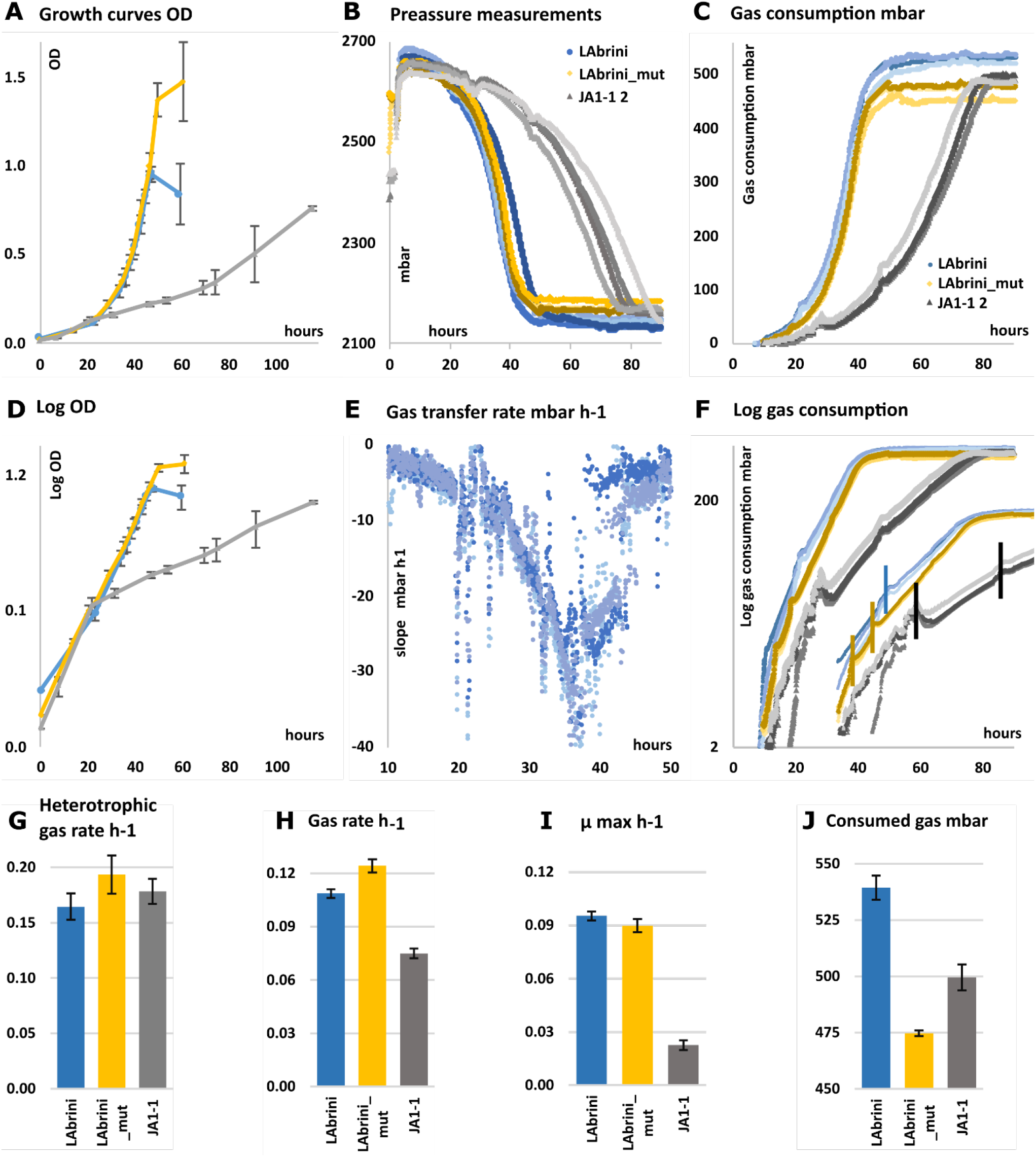
Pressure and OD characterisation of three *C. autoethanogenum* strains. A) OD growth curves with time (hours). B) Raw pressure reads (mbar) of the three strains. An initial pressure increase is observed as the temperature rises from room temperature to 37°C and as the gas changes from dry to wet due to water vapour pressure. C) Gas consumption of the three strains normalised to the highest pressure read for each individual vial. D) logarithmic plot of the OD samples with time (hours) E) Slope rate (mbar h-1) calculated at each sample time point (2 min), over a window of three samples, thus 6 minutes. F) Logarithmic plot of normalised gas consumption. The insert shows the identified exponential phases, separated by vertical lines. The correlation for each strain and the single cultivation is shown in supplementary figure 2,3,4. G) Gas consumption rates during the initial mixotrophic phase. H) Gas consumption rates h-1. For JA1-1 two exponential gas consumption phases were identified, here the fastest is shown between 34-44h. The second is 0.052 h-1 ±0.002 (53-67h). I) OD-based maximum growth rates observed (h-1). JA1-1 and LAbrini data from (Ingelman et al. 2024). J) Measured consumed gas mbar. Error bars are the standard deviation from 3 to 5 conditions. In pressure plots, all raw data points are shown, the curves are not fitted or smoothed.

We then compared the gas analysis and gas consumption rates with the OD obtained rates to assess the benefit of pressure sensors for this type of screening. The LAbrini and JA1-1 OD measurements are from (Ingelman et al. 2024). The OD measurements, and the pressure measurements are obtained from technical replicates and not the exact same experiment, as the OD sampling would introduce noise in the pressure data. Two of the strains, LAbrini (Ingelman et al. 2024) and LAbrini_mut, had similar biomass growth curves, and show statistically indifferent maximum specific growth rates (µ_max_) while JA1-1 grows a lot slower (Figure 1A,D). The growth rates for the three strains are quite similar until h 22, after which JA1-1 shifts to a slower biomass growth. This aligns with the initial faster, similar gas-uptake phase, hypothesised to be caused by the yeast extract, and the shift in the gas-uptake rate for JA1-1. In the biomass-based growth measurements, we thus can identify one of the shifts also seen in the gas-rate data. For LAbrini and LAbrini_mut, the catabolic shift from mixotrophic growth to pure-gas growth cannot be detected from the biomass data alone. For JA1-1 the gas consumption stops around 80h, while the biomass growth continues until 116h, thus 36 hours later. Across all cultivations, the time windows used to estimate exponential biomass growth were 10-15 hours later than those used for exponential gas consumption rates (Supplementary Figure 6). Taken together, this indicates potential growth delay due to OD sampling or insufficient OD data quality, though further verification is needed.

During the main autotrophic gas phase. Since cultivation relies solely on carbon and reducing power from CO, product rates can be estimated as the overall gas consumption rate minus the growth rate. LAbrini, LAbrini_mut, and JA1-1 have estimated product formation rates of 0.013, 0.034 and 0.052 h^-1^, respectively (Table 2). Thus, LAbrini_mut could have a 2.6 times higher product formation rate than Labrini, and JA1-1 could have 4 and 1.5 times higher estimated product rates than LAbrini and LAbrini_mut, respectively. Though only estimates, these are useful parameters for comparison during early screening. To further validate and obtain quantitative values, the correct biomass rates for each gas phase must be measured, along with the final gas composition and product concentrations, and the results verified against the elemental electron and carbon balances. For example, in this study, we cannot distinguish residual CO from residual or produced CO_2_. This is especially problematic for microbial communities or co-cultures, where parallel processes can consume and produce gas simultaneously (Borch, Grimalt-Alemany, et al. 2026). It is still a useful estimate and could, in some cases, provide physiological insights that other methods would lose. Manual OD sampling introduces disturbances and delays, a challenge that becomes increasingly problematic with slower-growing strains and smaller cultivation volumes. To validate the product estimates, online, non-intrusive measurements of biomass could be used. One option is to use backscatter. Depending on the cultivation goal, using online pressure measurements might be sufficient to achieve the desired characterisation and comparison.

**Table 2.**
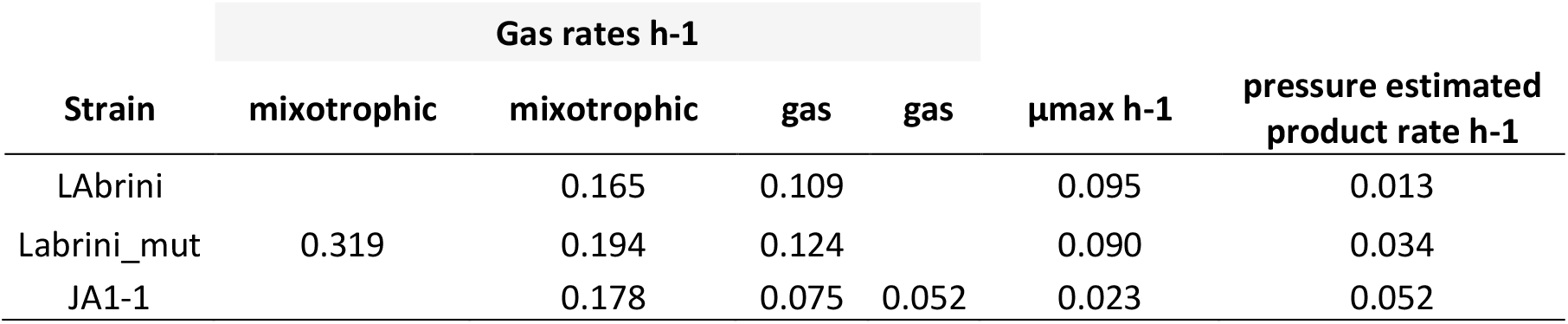
*Clostridium autoethanogenum* strain characteristics. Gas exchange rates are calculated from several exponential phases. OD-derived growth rates (µmax) are from the earlier experiment (Ingelman et al. 2024). Estimation of product rates, based on gas rates in the main gas-metabolic-phase minus the growth rate. This is not a qualitative molar calculation, but a screening comparison. Stochiometric balances, and simultaneous pressure and OD rates are needed for full calculation.

## 4 Discussion

### 4.1 Undisturbed final pressure thresholds and experimental design

The pressure-based data provides novel insights, some from the combination of a very high resolution and no sampling. The differences between total gas consumption and final pressures range from 43 to 65 mbar. This pressure difference could easily be caused by manual sampling alone. Every time the vials are sampled, some gas will be lost. Depending on the dead volume in the headspace of the sampling equipment and the pressure and volume of the cultivation vessel, the observed difference can be lost in one to four samples. Yet here, the difference is distinct and clear. Therefore, to accurately determine this strain difference, cultivation must proceed without sampling. The difference observed in the final pressure is an important parameter for strain characterisation. It corresponds to the lowest possible partial pressure of CO that supports the growth of the given bacteria (Philips 2020; Laura and Jo 2023). At an industrial scale, CO gas transfer is likely the rate-limiting step. Accurately measuring thresholds and selecting for microbes with low p(CO) thresholds in the early screening phase is important, as it could lead to improved growth performance under later, more limited conditions. This could have implications at an industrial scale.

One way to gain deeper insight into threshold performance is to cultivate individual strains or several strains under gas-limited conditions. The strain with the lowest threshold, and consequently the best gas-scavenging ability, will then flourish. Gas-limited conditions can be created by reducing shaking, thereby limiting gas transfer and establishing a gas-limited growth phase. This can be confirmed by the linear phase observed in the slope-scatter plot, as mentioned earlier. Sequential enrichment under these conditions serves as an adaptive evolution experiment for strains with the lowest gas thresholds. Under unconstrained growth with adequate gas transfer, as in this case, it also provides a means to select the fastest gas-consuming strains. Both approaches offer an alternative to the commonly used practice of selecting the fastest-growing strain, thus the fastest biomass formation.

This experiment had adequate gas transfer, as no linear phase in the gas consumption slope was observed. Under the conditions used, the exponential growth phase in the gas could be prolonged, as growth is not yet limited by gas transfer. To achieve this, the CO content in the batch cultivations needs to be increased. Since the final pressure is very high, some of the residual N_2_ or CO_2_ could be replaced by CO during the experimental preparation. The initial, faster phase of mixotrophic gas consumption could be reduced or eliminated by limiting yeast extract in the medium.

### 4.2 Online pressure and novelty

Pressure or CO_2_-based cultivation measurements are not new in microbiology, and the pressure response in closed systems has been extensively discussed for gas uptake and production rates, also in relation to the solubility and buffering effect of CO_2_ (Myers and Matsen 1955). It is a common measurement in biodegradation, soil-respiration, food-, rumen-, and gas-fermentation, inter-species interaction, and for analysing physiological response due to blood pressure cycles (Freijer and Bouten 1991; Rotaru et al. 2021; Szmelter et al. 2022; Laura and Jo 2023; Bond-Lamberty et al. 2024; Liu et al. 2025; Pérez-Vidal et al. 2025). In anaerobic batch cultures, the pressure is often sampled manually, and as single endpoint measurements, but tools with online pressure are available for these analyses, particularly used for biogas, or bio-methanation potential (BMP) (Hofmann et al. 2025). The prior art underscores the importance of pressure effects on biological metabolism and the diversity of insights that can be gained from high-quality pressure data.

The key aspects and novelty of the present study, and the tools and methodology presented here, lie in the simultaneous combination and focus on several parameters. It is high-quality data gathered from undisturbed, high-resolution, real-time pressure measurements in small-scale closed anaerobic sterile batch cultivations. This enabled the detection of metabolic shifts through changes in the catabolic rate, which was not identified through OD-based growth rate measurements alone. Additionally, the method identifies the gas thresholds at which the driving forces for the involved catabolic reactions reach the minimum energy required for growth.

## 5 Conclusion and perspectives

Overall, the online gas pressure measurements enabled evaluation of the growth setup, characterised it, compared it, and separated the strains based on the detection of several metabolic shifts in gas consumption rate and final pressure. These insights had not been previously identified from the OD measurements alone. The work establishes a practical framework for catabolism-resolved microbial characterisation in sealed batch vials through high-resolution online pressure (gas exchange). The assumption that pressure measurements correlate with CO_2_ and catabolic rates is sensitive to solubility/buffering and temperature/vapour effects, but these limitations are addressable through controls and complementary analytics. At the same time, the pressure measurement and a simple design enable a robust, easy-to-use method that integrates well with existing sterile anaerobic cultivation workflows. It is particularly suitable for cultivation systems where optical measurements cannot be applied, and for measuring non- or slowly growing anaerobic cultures and mixed communities, where substantial activity may occur through interaction cascades with limited changes in total biomass. While similar analyses can be performed in large, instrumented bioreactors, the advantage here is a small-scale, low-cost, no-headspace-disturbance setup that allows such characterisation early in a strain selection and community development process, reducing reliance solely on biomass curves for conclusions. Also, the online measurements and a design and form factor that integrate with existing bottle- or vial-based cultivations allow for easy scaling of the conditions tested.

## Supporting information

mmborch 2026 - Measure Catabolism - supplementary file

## 6 Acknowledgments

The first author is grateful for the collaboration and appreciates the warm welcome in Estonia and the great conversations that hopefully will continue.

## 7 Declarations

### Author Contributions and Information

Conceptualisation, Methodology, hardware development, project administration, writing – original draft, MMB. Investigation, formal analysis: MMB, UJN. Writing – review & editing: ATN, MMB, PGV, KV, UJN. Supervision, funding acquisition: ATN, KV.

### Data Availability Statement

The datasets generated and analysed for this study can be made available upon reasonable request.

