## Supplementary material for "Measure Catabolism: Real-time shifts in microbial metabolism through online pressure measurements": mmborch 2026 - Measure Catabolism - supplementary file

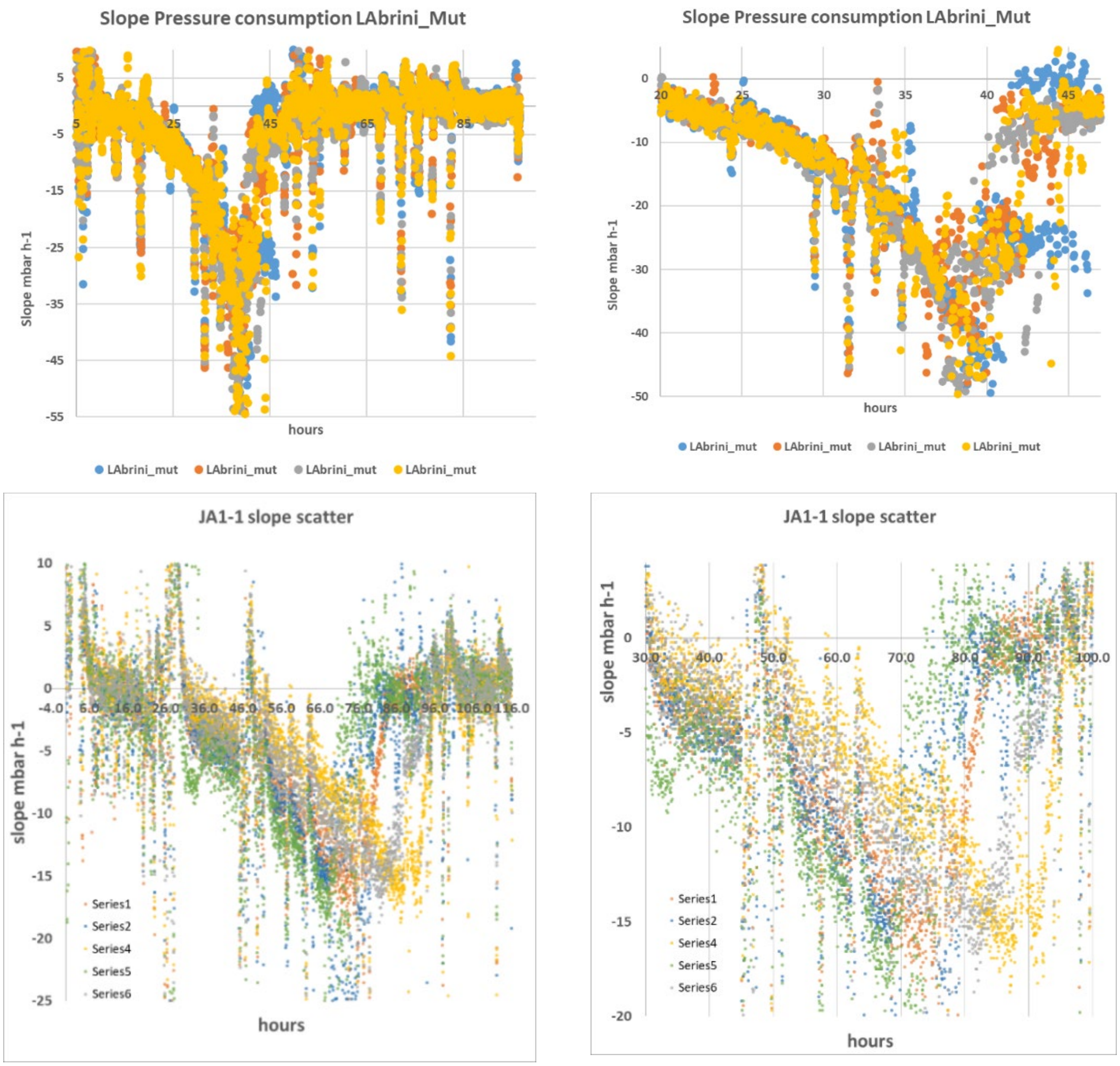


Supplementary Figure 1. A) Slope of LAbrini_mut and zoom B) slope scatter of JA1-1 and zoom

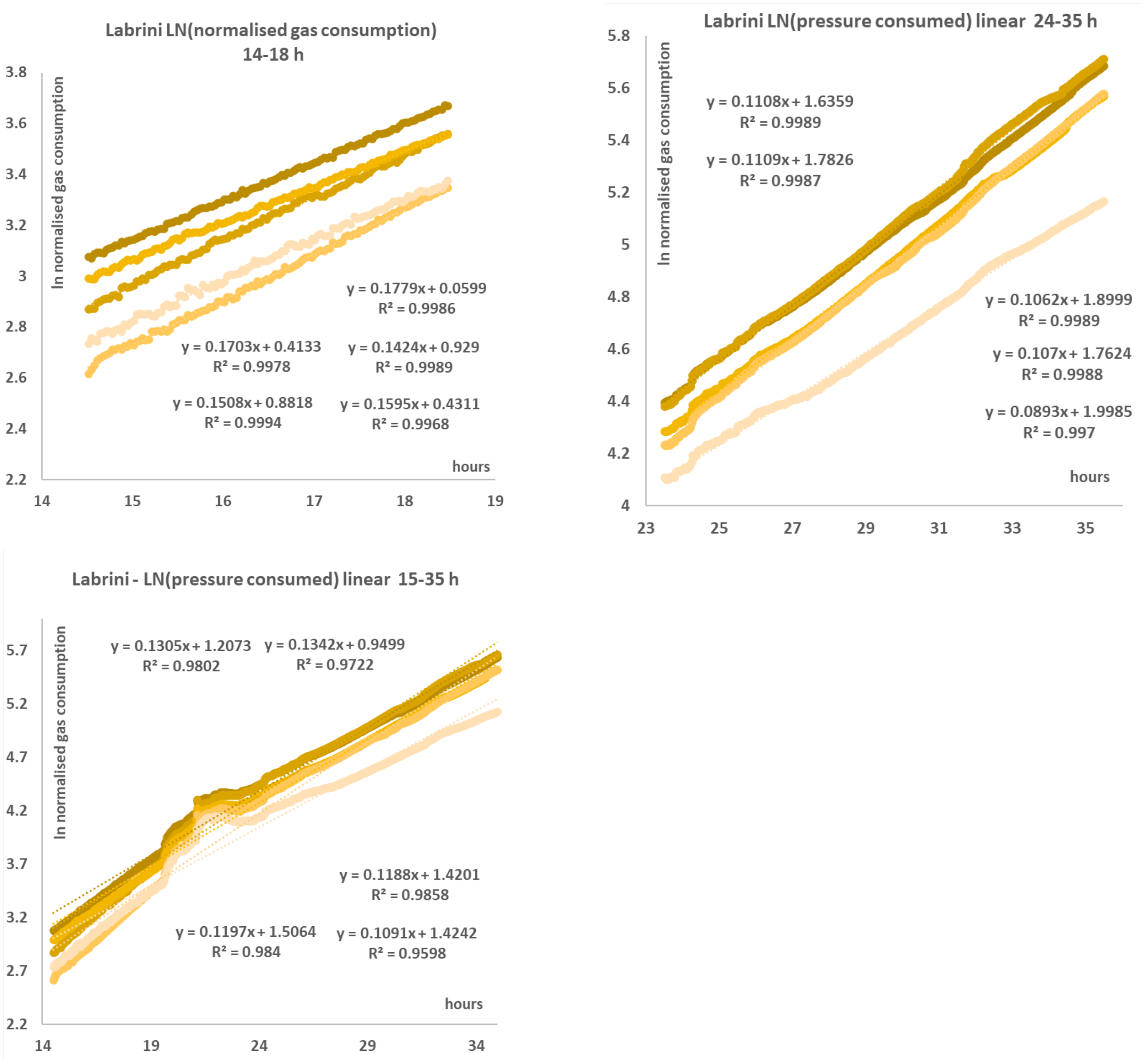


Supplementary Figure 2. Gas consumption rates of LAbrini from linear fit of ln(normalised gas consumption). A) Early fit h 14-18. B) fit during h 24 – 35. C) Fit during the two phases combined, h 15-35.

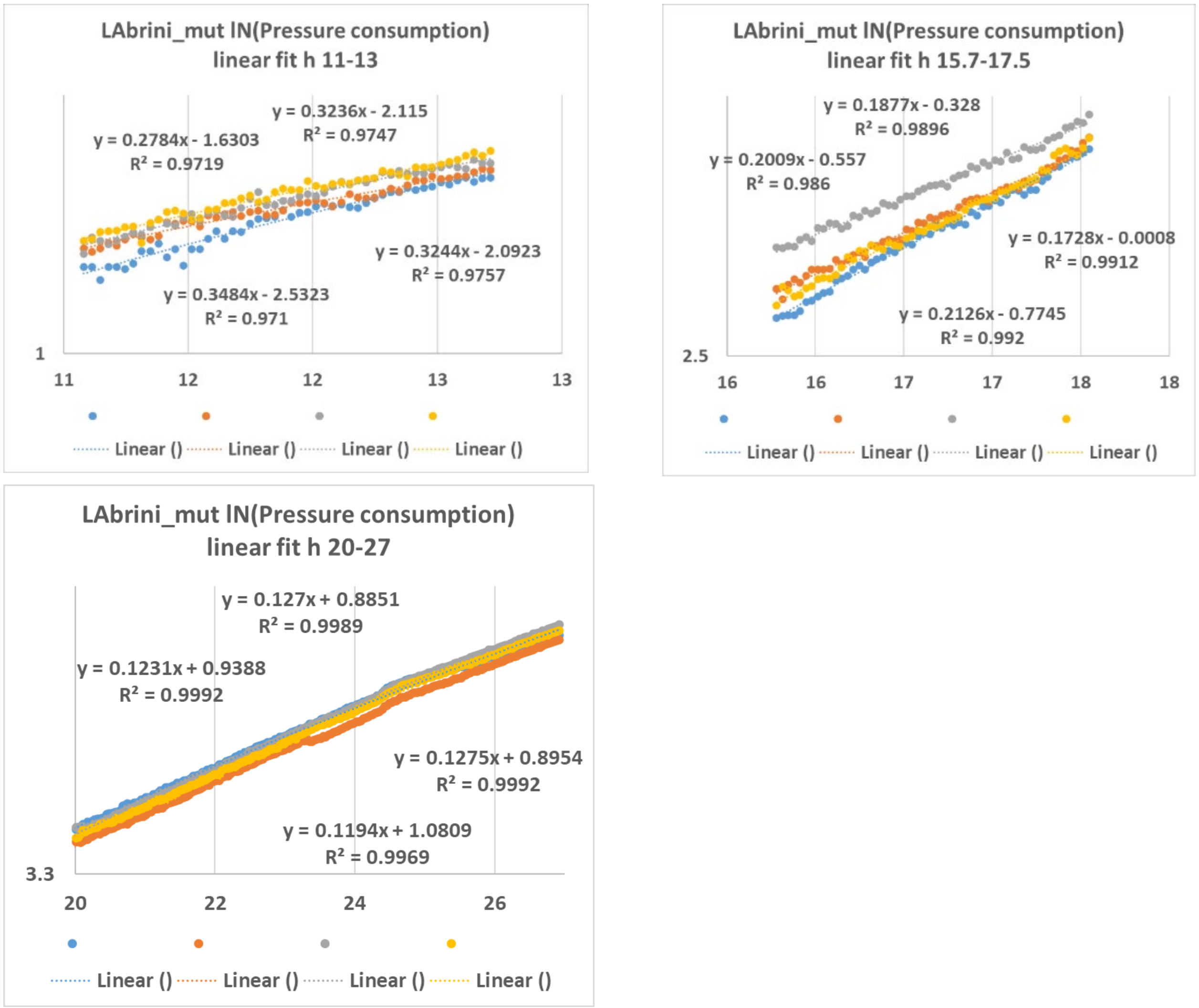


Supplementary Figure 3. Gas consumption rates of LAbrini_mut from linear fit of ln( normalised gas consumption) to investigate the three phases identified. A) early fit h 11-13. B) fit during h 15.7 – 17.5. C) fit during gas growth only, h 20-27.

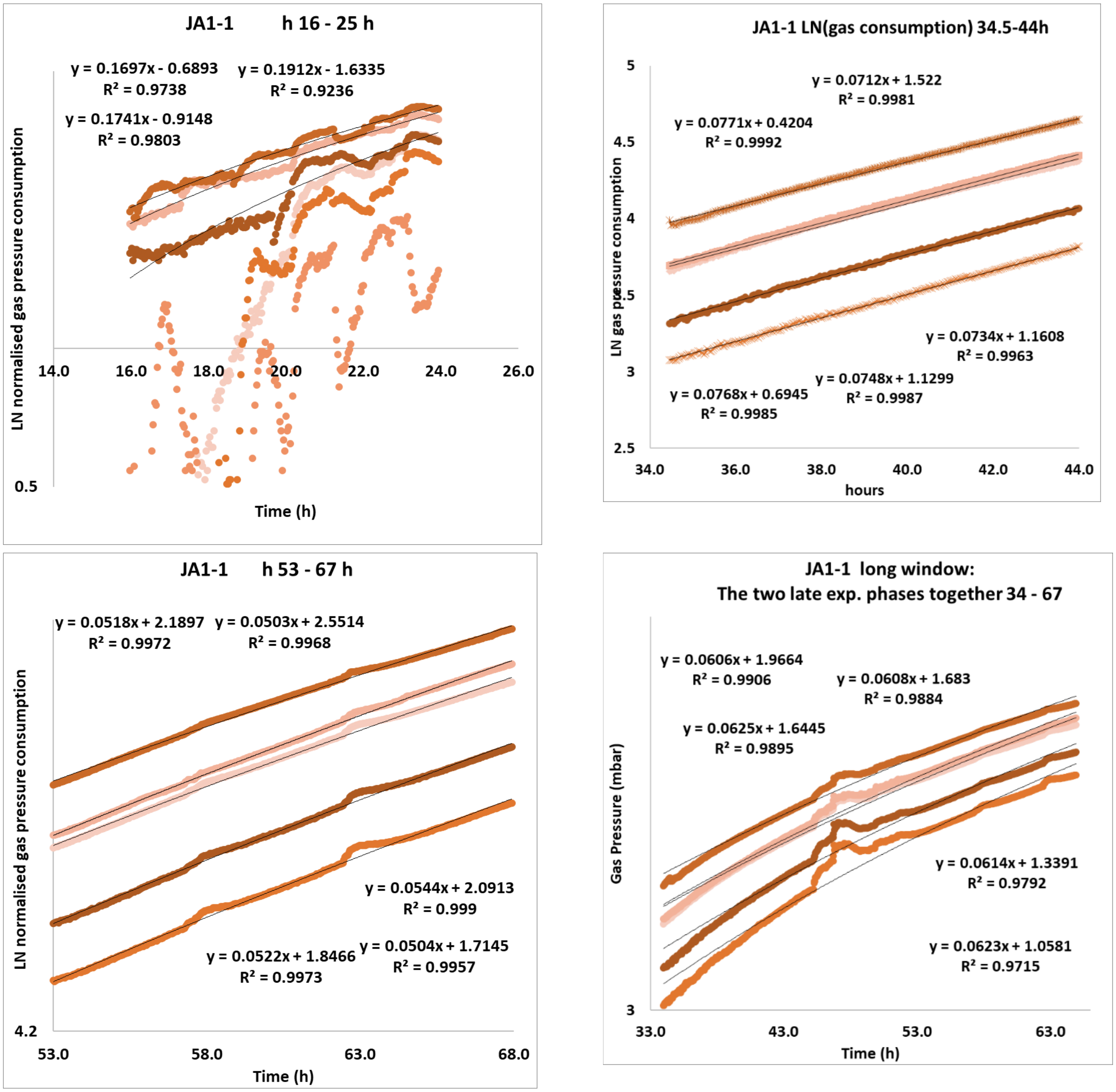


Supplementary Figure 4. Gas consumption rate fitting of the strain JA1-1, for different time windows, to verify exponential phases. Linear fit to ln(normalised gas consumption). A) h 16 – 25, B) h 34.5 – 44. C) h 54-67 and D) h 34 – 67.

| Gas rates mmbar h-1 | LAbrini | | | LAbrini_ mut | | | JA1-1 | | |
|---|---|---|---|---|---|---|---|---|---|
| "Phase" | 1 | 2 | 3 | 1 | 2 | 3 | 1 | 2 | 3 |
| hours | | 14-18h | 24-34 h | 11-13 | 15.7-17.5 | 20-30 | 15 - 25 h | 34.5-44h | 53 - 67 h |
| Mean Rate | | 0.165 | 0.109 | 0.319 | 0.194 | 0.124 | 0.178 | 0.075 | 0.052 |
| SD | | 0.012 | 0.002 | 0.029 | 0.017 | 0.004 | 0.011 | 0.002 | 0.002 |
| rep 1 | | 0.178 | 0.111 | 0.278 | 0.188 | 0.123 | 0.174 | 0.077 | 0.050 |
| rep 2 | | 0.170 | 0.111 | 0.324 | 0.201 | 0.127 | 0.170 | 0.071 | 0.052 |
| rep 3 | | 0.151 | 0.106 | 0.348 | 0.213 | 0.128 | 0.191 | 0.073 | 0.054 |
| rep 4 | | 0.160 | 0.107 | 0.324 | 0.173 | 0.119 | | 0.075 | 0.052 |
| rep 5 | | | | | | | | 0.077 | 0.050 |

Supplementary Table 1. Gas rate from the identified exponential gas consumption phases

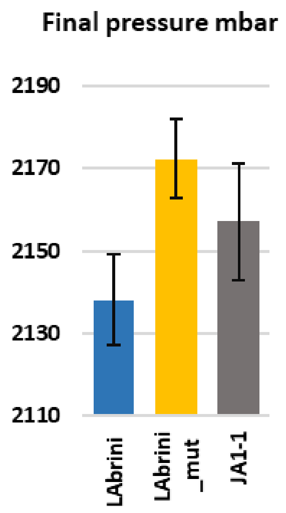


Supplementary Figure 5. Final pressure mbar.

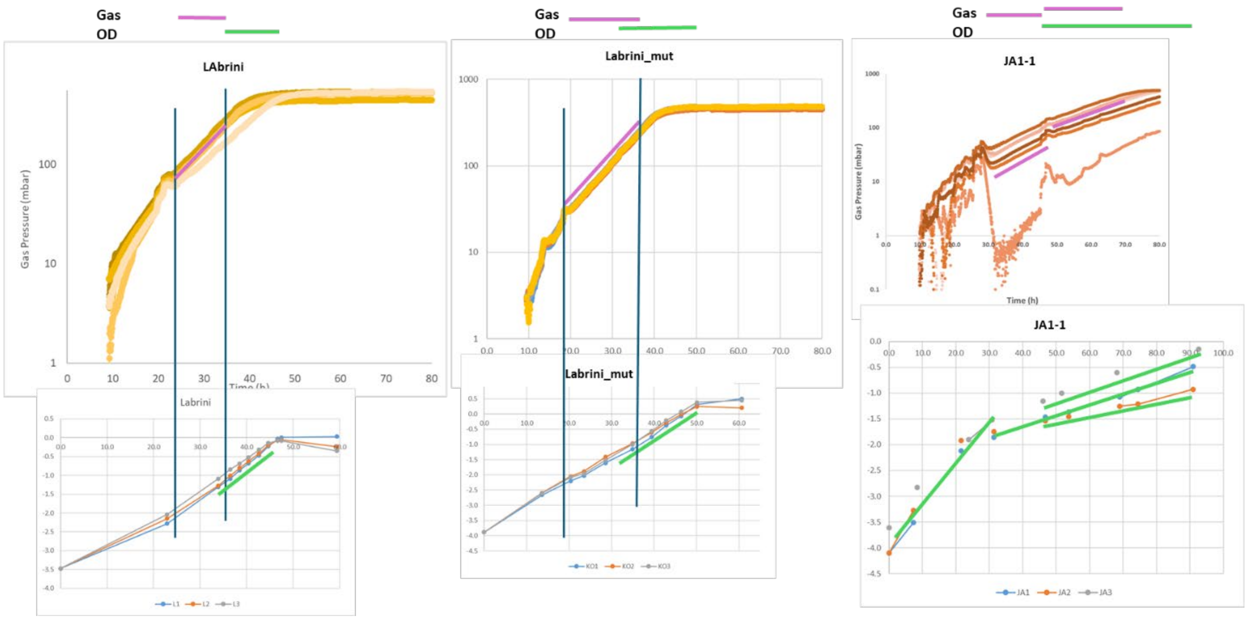


**Supplementary Figure 6.** Comparison of the time window for exponential growth and gas uptake rates. Data from separate experiments. Manual sampling appears to postpone the time window 10-15h.
